# Effects of Gene Dosage and Development on Subcortical Nuclei Volumes in Individuals with 22q11.2 Copy Number Variations

**DOI:** 10.1101/2023.10.31.564553

**Authors:** Charles H. Schleifer, Kathleen P. O’Hora, Hoki Fung, Jennifer Xu, Taylor-Ann Robinson, Angela S. Wu, Leila Kushan-Wells, Amy Lin, Christopher R. K. Ching, Carrie E. Bearden

## Abstract

The 22q11.2 locus contains genes critical for brain development. Reciprocal Copy Number Variations (CNVs) at this locus impact risk for neurodevelopmental and psychiatric disorders. Both 22q11.2 deletions (22qDel) and duplications (22qDup) are associated with autism, but 22qDel uniquely elevates schizophrenia risk. Understanding brain phenotypes associated with these highly penetrant CNVs can provide insights into genetic pathways underlying neuropsychiatric disorders. Human neuroimaging and animal models indicate subcortical brain alterations in 22qDel, yet little is known about developmental differences across specific nuclei between reciprocal 22q11.2 CNV carriers and typically developing (TD) controls. We conducted a longitudinal MRI study in 22qDel (n=96, 53.1% female), 22qDup (n=37, 45.9% female), and TD controls (n=80, 51.2% female), across a wide age range (5.5-49.5 years). Volumes of the thalamus, hippocampus, amygdala, and anatomical subregions were estimated using FreeSurfer, and the effect of 22q11.2 gene dosage was examined using linear mixed models. Age-related changes were characterized with general additive mixed models (GAMMs). Positive gene dosage effects (22qDel < TD < 22qDup) were observed for total intracranial and whole hippocampus volumes, but not whole thalamus or amygdala volumes. Several amygdala subregions exhibited similar positive effects, with bi-directional effects found across thalamic nuclei. Distinct age-related trajectories were observed across the three groups. Notably, both 22qDel and 22qDup carriers exhibited flattened development of hippocampal CA2/3 subfields relative to TD controls. This study provides novel insights into the impact of 22q11.2 CNVs on subcortical brain structures and their developmental trajectories.

## Introduction

Genomic copy number variations (CNVs) at the 22q11.2 locus strongly increase risk for neurodevelopmental and psychiatric disorders including autism and schizophrenia [1]. 22q11.2 Deletion Syndrome (22qDel) and 22q11.2 Duplication Syndrome (22qDup) result from reciprocal CNVs that involve hemizygous deletion or duplication of approximately 2.6 Megabases (Mb) of genomic material from the long arm of chromosome 22. The brain and behavioral phenotypes resulting from these related CNVs provide a valuable genetics-first framework for investigating biological pathways relevant to brain development and neuropsychiatric disorders [2,3].

22qDel (OMIM #188400, #192430) is one of the strongest known genetic risk factors for schizophrenia, with over 1 in 10 individuals with 22qDel having a comorbid psychotic disorder and over one-third experiencing subthreshold psychosis symptoms [4,5]. 22qDel also increases risk for autism, attention-deficit hyperactivity disorder (ADHD), intellectual disability, and anxiety disorders [6–8]. This microdeletion occurs in approximately 1 in 4000 people [9].

A duplication of this same region causes 22qDup (OMIM #608363) and is often inherited, unlike 22qDel which typically arises *de novo* [10]. 22qDup was discovered more recently than 22qDel and has not yet been as deeply characterized [11,12]. Individuals with 22qDup experience higher rates of neurodevelopmental disorders, including intellectual disability and autism, compared to the general population; however, the duplication generally has a milder impact on neurodevelopment compared to 22qDel [1,13]. In contrast to 22qDel, 22qDup is *less* common in individuals with schizophrenia compared to the general population, suggesting a potential protective effect against schizophrenia in 22qDup [14–16].

In addition to widespread cortical anomalies, including reductions in surface area, concomitant with relatively increased cortical thickness [17,18], studies of 22qDel have consistently identified structural and functional alterations in subcortical structures. A large multi-site, cross-sectional study from the ENIGMA 22q11.2 Working Group found decreased subcortical volumes in 22qDel compared to TD controls, with larger effects in those with psychosis [19]. These subcortical brain structures play key roles in cognitive, sensory, and affective processes [20]. Individual differences in subcortical anatomy have been related to both common and rare genetic variation [21,22], and to psychiatric and neurodevelopmental disorders including schizophrenia, autism, and ADHD [23,24].

While most of the literature to date has investigated whole subcortical structures and/or voxel-wise shape differences, these structures are composed of many small distinct nuclei [20,25]. More recent studies have begun to investigate volumes of specific anatomical subregions of structures such as the thalamus and hippocampus. Bi-directional effects on FreeSurfer-derived thalamic nuclei volumes have been observed in 22qDel, wherein volumes of subregions involved in sensory processes (e.g., medial geniculate) were found to be smaller than controls, while subregions involved in cognitive processes (e.g., anteroventral) were larger [26]. Longitudinally, there were steeper thalamic volume decreases over time in individuals with 22qDel who experienced auditory hallucinations. In 22qDel, lower hippocampal tail volume has been related to verbal learning impairments [27], and hippocampal volume loss over time in 22qDel has been linked to altered local balance of excitatory and inhibitory neurotransmitter metabolites [28].

Few studies have directly compared brain phenotypes in 22qDel and 22qDup. In the first study comparing regional brain volumes of reciprocal 22q11.2 CNV carriers to typically developing (TD) controls, in a cross-sectional sample our group found that gene dosage (i.e., the number of copies of the 22q11.2 locus) was positively related to cortical surface area (22qDel < TD < 22qDup) and negatively related to cortical thickness (22qDel > TD > 22qDup) [18]. This study also found larger hippocampal volumes in 22qDup relative to 22qDel, and radial thickness differences in subcortical structures, including the thalamus and amygdala. Recently, a large study of subcortical volumes in 11 different CNVs found convergent evidence for hippocampal volume differences in 22qDel versus 22qDup [21].

No study has assessed subcortical subregional volumes or longitudinal subcortical development in 22qDup, nor directly compared subregion-level volumes between 22qDup and 22qDel. Studies of subcortical shape show complex alterations in 22q11.2 CNV carriers compared to controls, with localized volume increases and decreases suggesting differential vulnerabilities across subregions [19,21]. Mapping gene dosage effects to functionally and histologically defined nuclei can facilitate causal links between macro-scale MRI brain signatures and cellular and molecular mechanisms inferred from other data sources including post-mortem brain tissue [29,30], and animal models [31]. Examples of genes in the 22q11.2 locus that have been related to cortical structural phenotypes and which may have broad or region-specific effects on subcortical development include *DGCR8*, a gene involved in microRNA regulation, and *AIFM3*, a gene involved in apoptosis pathways [32].

In this longitudinal structural MRI study of reciprocal 22q11.2 CNV carriers and TD controls, across a wide age range (ages 5.5-49.5), we present the first investigation of the effects of gene dosage at the 22q11.2 locus on anatomical subregion volumes in the thalamus, hippocampus, and amygdala. We also characterize, for the first time, developmental trajectories of these subcortical volumes in individuals with 22q11.2 CNVs.

## Methods

### Participants

The total longitudinal sample consisted of 387 scans from 213 participants (5.5–49.5 years of age; n=96 22qDel baseline; n=37 22qDup baseline; n=80 TD controls baseline; see **Table 1**, and **Figure S1**). Participants had data from 1-6 timepoints separated by an average of approximately 1.75 years. The groups were matched on baseline age and sex, mean number of longitudinal visits, and interval between visits. See **Supplemental Methods** for details on inclusion/exclusion criteria and clinical assessment. After study procedures had been fully explained, adult participants provided written consent, while participants under the age of 18 years provided written assent with the written consent of their parent/guardian. The UCLA Institutional Review Board approved all study procedures and informed consent documents.

**Table 1.**
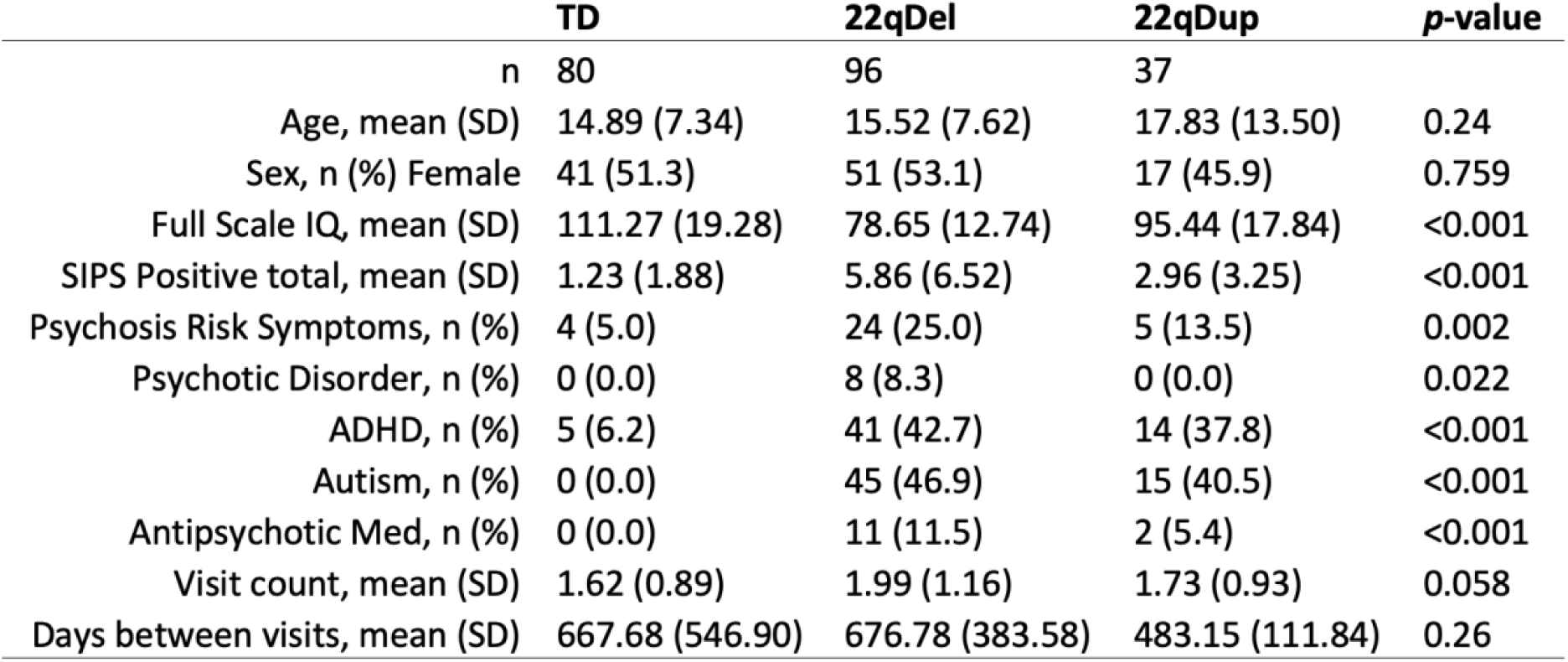
Baseline Demographics. TD controls, 22qDel, and 22qDup with *p*-values for between group comparisons (ANOVA for continuous variables and chi-squared for categorical). Baseline cohorts are statistically matched based on age and sex as well as mean number of longitudinal visits and interval between visits. Cognition was measured with the Wechsler Abbreviated Scale of Intelligence-2 (WASI-2). Prodromal (psychosis-risk) symptoms were assessed with the Structured Interview for Psychosis-Risk Syndromes (SIPS). Psychosis Risk Symptoms are operationalized here as having any score of 3 or greater (i.e., prodromal range) on any SIPS positive symptom item. Psychotic disorder diagnosis is based on structured clinical interview (SCID) for DSM-IV/V and includes schizophrenia, schizoaffective disorder, brief psychotic disorder, and psychotic disorder not otherwise specified.

#### Neuroimaging acquisition/preprocessing

All participants were imaged at the UCLA Center for Cognitive Neuroscience on either a Siemens TimTrio or Siemens Prisma scanner with the same T1-weighted (T1w) sequence [33]. Scan sessions at all timepoints were first processed cross-sectionally using the recon-all anatomical segmentation pipeline in FreeSurfer 7.3.2 [34–36]. The FreeSurfer longitudinal stream was subsequently applied, which has been shown to significantly improve reliability and statistical power in repeated measure analyses [37].

#### Subcortical nuclei

For each scan, volume was estimated for the whole thalamus, amygdala, and hippocampus, and their subregions using Bayesian methods to automatically segment T1w images using template atlases based on histological data and ultra-high-resolution *ex vivo* MRI [38–41]. These segmentations are well-validated and have been applied by multiple consortia to large scale neuroimaging analyses [42–46]. See **Supplemental Methods** for details on segmentation, qualitative and statistical quality control (QC) procedures, and between-scanner harmonization using longitudinal ComBat [47].

#### Gene dosage effects

To investigate the impact of CNV status on subcortical volumes, we used a linear mixed effects approach where gene dosage was numerically coded based on CNV status: 22qDel=1, TD=2, and 22qDup=3 copies of the 22q11.2 locus. First, total intracranial volume (ICV) was normalized based on the TD group mean and standard deviation, and a random effects model was tested predicting total ICV from gene dosage, controlling for linear and quadratic age, sex, and scanner, with a random intercept for each participant. Next, for each subcortical region, ComBat-adjusted volumes were normalized, and a random effects model was tested with the same covariates plus ICV. Gene dosage effects were tested for the whole thalamus, hippocampus, and amygdala volumes, followed by each of the 40 subregions. All tests were corrected for multiple comparisons using the standard False Discovery Rate (FDR) at a threshold of *q*<0.05 across the 44 volumes [48].

#### Maturational effects

To characterize developmental trajectories, we assessed nonlinear age-related changes with general additive mixed models (GAMMs) as in Jalbrzikowski et al., 2022 [49]. GAMMs are a nonlinear extension of mixed effects regression and are well-suited to data with repeated measures [50]. For each subcortical volume, a GAMM was computed predicting volume from age in each group, controlling for sex, ICV, and scanner, with a random intercept for each participant. p-values for the effect of age in each group were FDR-corrected across all 129 models. Age ranges of significant difference between CNV groups and controls were computed from the 95% confidence interval for the difference in curves.

#### Secondary analyses

Regional volume differences compared to the TD group were tested separately for 22qDel and 22qDup groups. Gene dosage analyses were also repeated without averaging structures bilaterally, to detect any asymmetric hemispheric effects. Secondary analyses of the interaction between sex and gene dosage on brain volumes were also tested. The effect of antipsychotic medication was also investigated.

Motivated by literature relating low hippocampal tail volume to verbal learning impairment in 22qDel [27], we assessed verbal and non-verbal IQ [Wechsler Abbreviated Scale of Intelligence (WASI-2) Vocabulary and Matrix subtest scaled scores] for associations with hippocampal tail volume in each group. Given the relationship between 22qDel and psychosis risk [9,26,51], we additionally tested relationships between psychosis risk symptoms and all subcortical volumes in the 22qDel group.

## Results

### Gene dosage effects

Total ICV was positively related to gene dosage (**Table 2**, **Figure 1**). Positive gene dosage effects on volume were observed for the whole hippocampus, but not the whole thalamus or amygdala.

**Figure 1.**
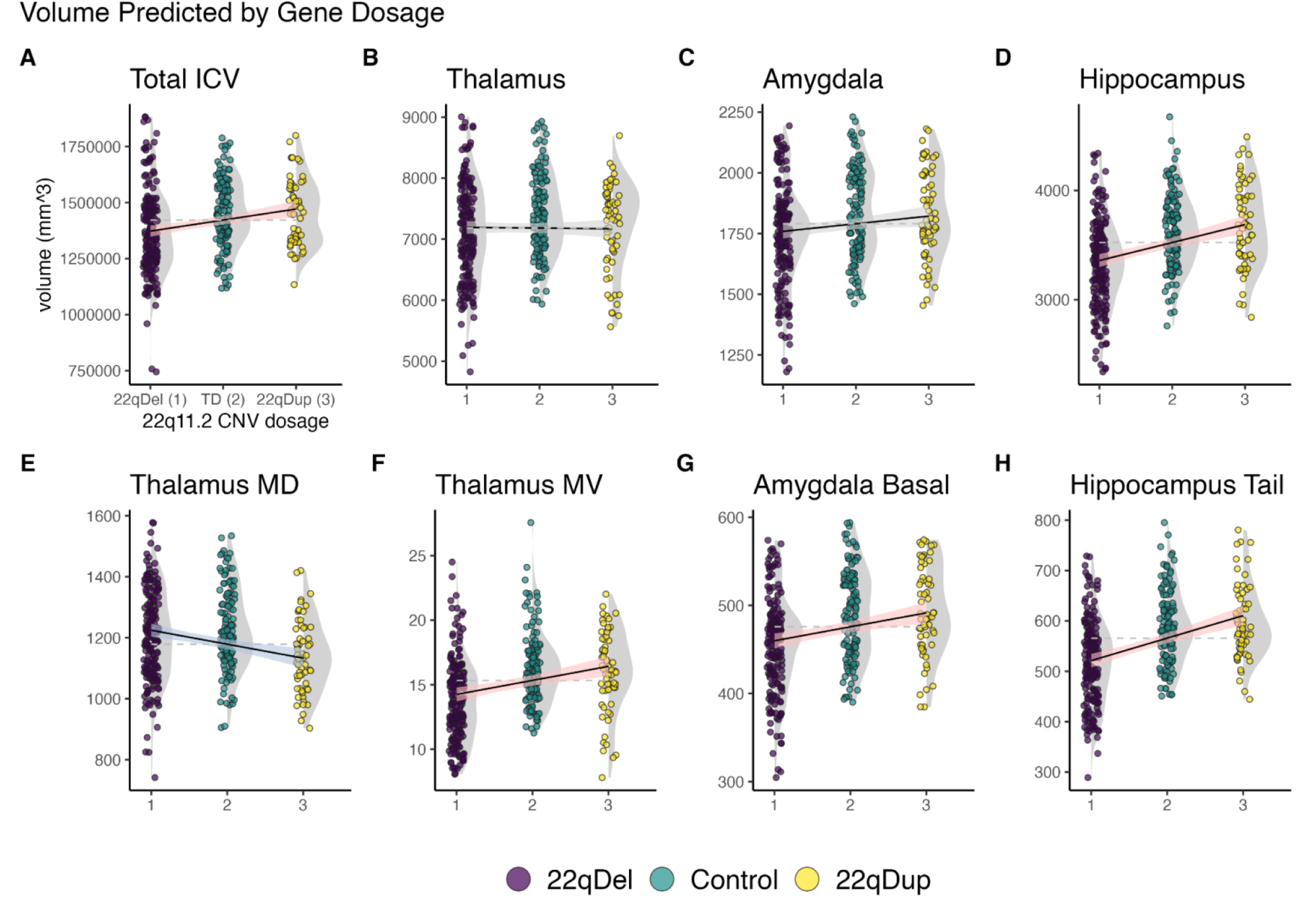
Visualization of gene dosage effects on selected volumes. Volume scatterplots for 22qDel (purple) control (green) and 22qDup (yellow), with kernel density estimates for each group in gray, overlaid with best fit lines for linear mixed models showing relationships between CNV dosage and regional volumes, with 95% confidence intervals (CI). Red CI indicates significant positive gene dosage relationship (FDR *q* < 0.05), blue indicates significant negative, and gray indicates no relationship. **A)** Total intracranial volume (ICV). All other models control for ICV. **B-D)** There was a significant positive relationship between gene dosage and whole hippocampus volume, which was not observed for the thalamus or amygdala. **E-F)** Within the thalamus, some subregions (e.g., MD; mediodorsal) showed negative gene dosage volume relationships, while others (e.g., MV; medial ventral) exhibited positive effects. **G)** Several amygdala subregions (e.g., basal nucleus) exhibited positive gene dosage effects, despite the whole amygdala having no relationship between gene dosage and volume. **H)** Significant hippocampal subregions (e.g., hippocampal tail) had the same direction of effect as the whole hippocampus.

**Table 2.**
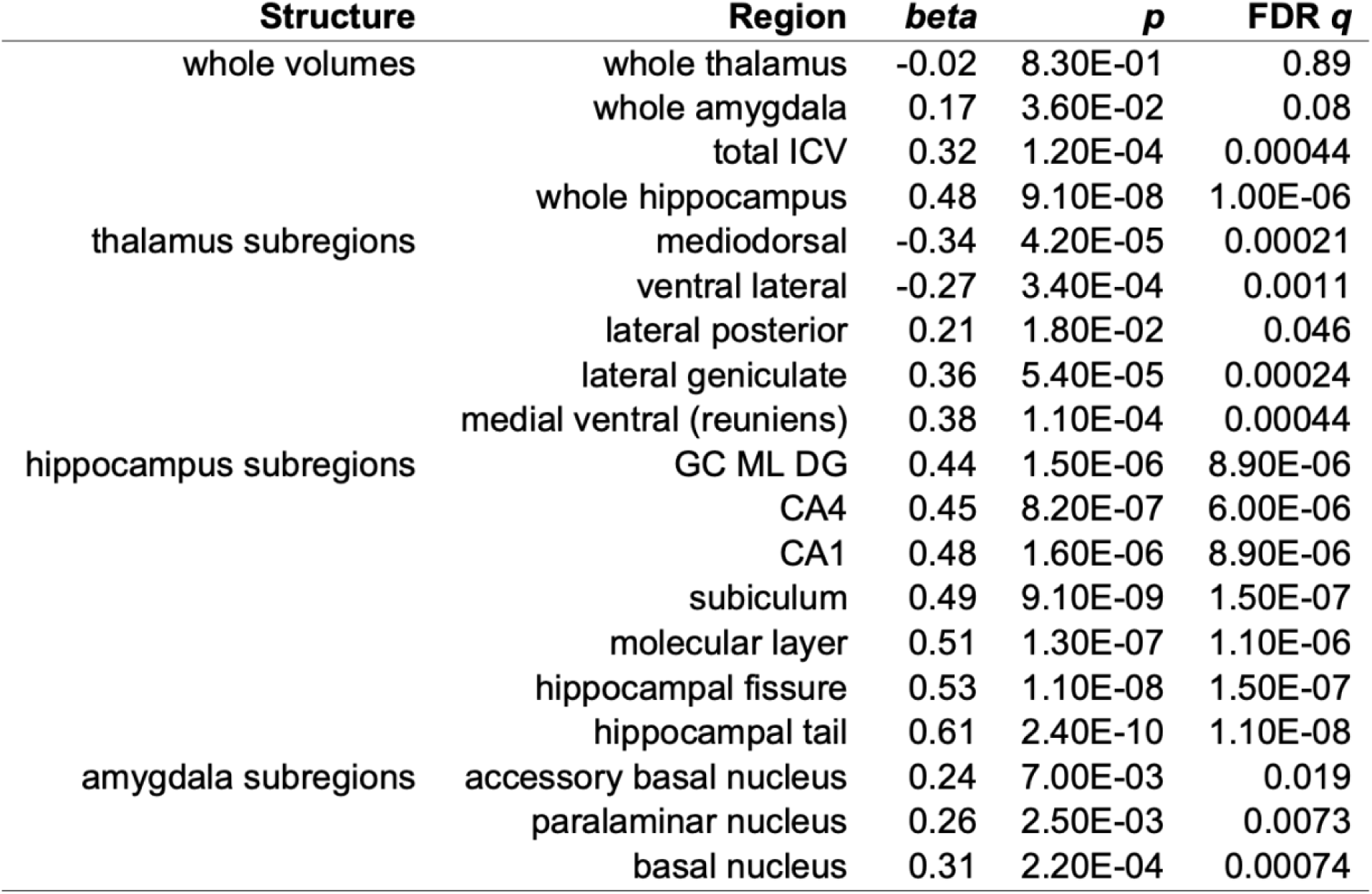
Gene dosage effects on subcortical volumes. Linear mixed models predicting normalized brain volumes from gene dosage (22qDel=1, TD=2, 22qDup=3). Gene dosage was positively related to total intracranial volume and whole hippocampus volume, but not to the whole thalamus or amygdala. Bi-directional effects within the thalamus, and localized amygdala effects were observable at the subregion level. Gene dosage was negatively related to volume for thalamic mediodorsal and ventral lateral regions. All other subregions of the thalamus, hippocampus, or amygdala with significant effects exhibited positive relationships between gene dosage and volume. Results are presented for all whole structure volumes (whole thalamus, amygdala, hippocampus, and total intracranial volume), and for subregions with significant False Discovery Rate (FDR) corrected gene dosage effects (FDR *q* < 0.05). Abbreviations: ICV, intracranial volume; GC ML DG, granule cell and molecular layer of the dentate gyrus; CA, cornu ammonis.

Within the thalamus and amygdala, individual subregions showed significant gene dosage effects (**Table 2**, **Figure 1**). There were bi-directional gene dosage effects on thalamic volumes; negative gene dosage effects were observed for the mediodorsal and ventral lateral nuclei, while positive gene dosage effects were observed in the lateral posterior, lateral geniculate, and reuniens nuclei. Within the amygdala, positive gene dosage effects were observed for the accessory basal, paralaminar, and basal nuclei. Multiple hippocampal subregions exhibited positive gene dosage effects in line with the whole structure findings, with the strongest effect in the hippocampal tail.

### Maturational effects

GAMM analysis revealed multiple subcortical regions with significant age-related changes in each cohort (**Figures 2&3** and **Table S4**). Overall, 22qDel and TD cohorts exhibited age-related changes across various subregions in the thalamus, hippocampus, and amygdala, while in 22qDup, age-related changes were only detected in thalamic regions. All three groups exhibited significant age-related decreases in thalamic mediodorsal and medial geniculate volumes, but medial geniculate volumes in 22qDup increased slightly after approximately age 30, whereas in the other groups they continued to decrease. Developmental trajectories after age 30 should be interpreted with caution, though, due to the lower number of participants in this age range.

**Figure 2.**
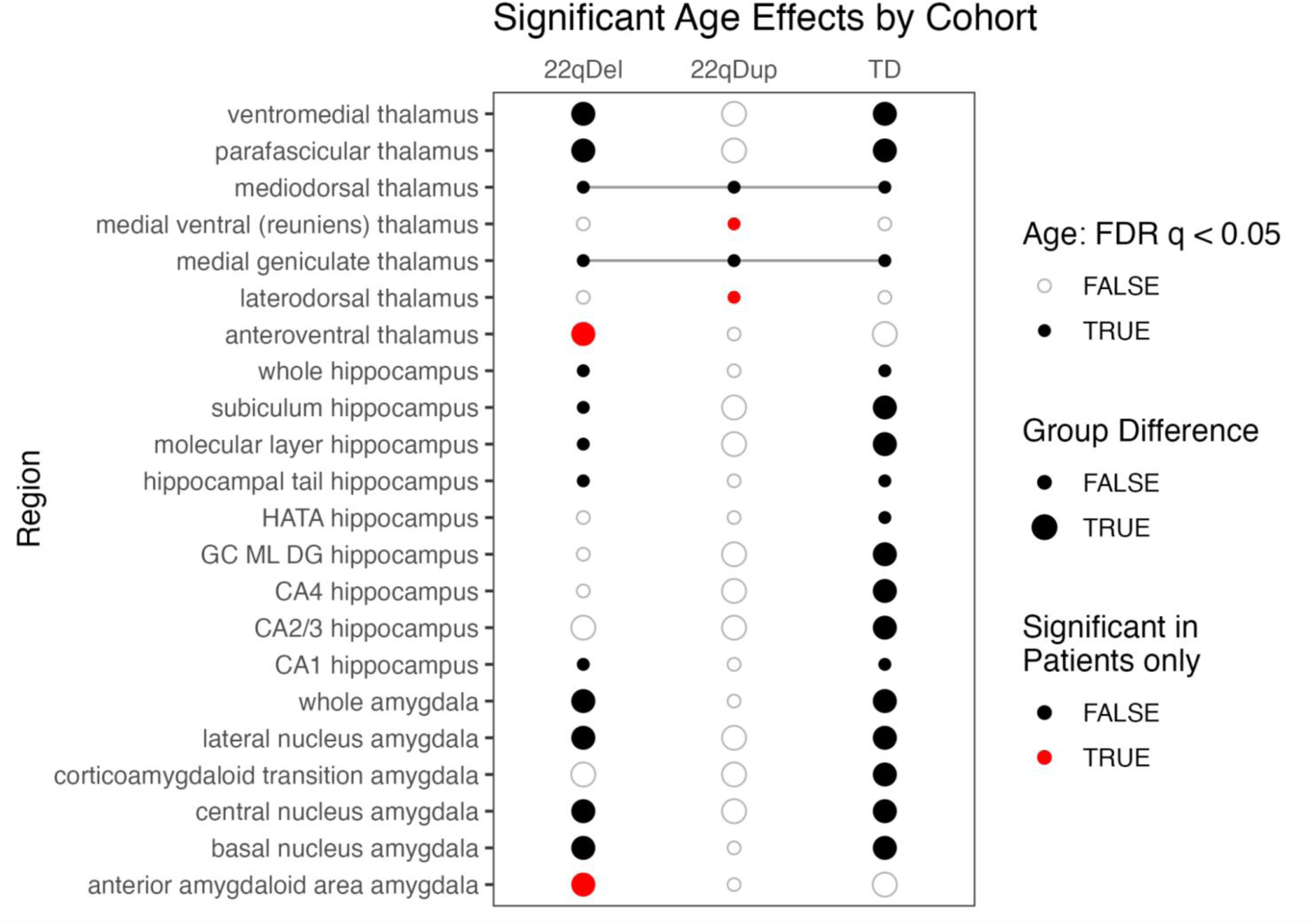
Summary of age effects on subcortical volumes. For each group, regions with FDR- corrected significant age effects on volume (*q* < 0.05) are marked with a dark circle. Large circles indicate at least one age range with a significant difference between CNV patients (22qDel or 22qDup) and TD control age curves based on the 95% confidence interval (CI), whereas small circles indicate overlapping patient and control curves. Red circles indicate regions that are significant in either patient group but not controls. Lines connect regions with significant age effects in all three groups.

Several regions exhibited significant age-related changes in only CNV carriers, but not TD controls (**Figure 2**). The anteroventral thalamus showed age effects in only 22qDel, involving steeper decreases in childhood and adolescence compared to the other groups (**Figure 3**). 22qDel volumes were significantly higher than TD between ages 5.5-8.7, but lower between ages 14.7-22.9 (**Table S4**). The anterior amygdaloid area also showed significant age effects in only 22qDel, but the curves for the three groups overlapped until age 35.5. Medial ventral and laterodorsal thalamus showed significant age effects in only 22qDup, but the curves overlapped TD at all ages. Several hippocampal subregions and the cotricoamygdaloid transition area exhibited significant age effects in TD controls but neither CNV group. In the hippocampal CA2/3 and CA4 regions, TD controls exhibited an inverted-U-shaped developmental curve, which was mostly flattened in 22qDel and 22qDup (**Figure 3**). CA2/3 volumes in 22qDel were greater than TD between ages 5.5-8.3 and lower than TD between ages 13.4-21.2, with similar periods of difference for 22qDup versus TD (**Table S4**).

**Figure 3.**
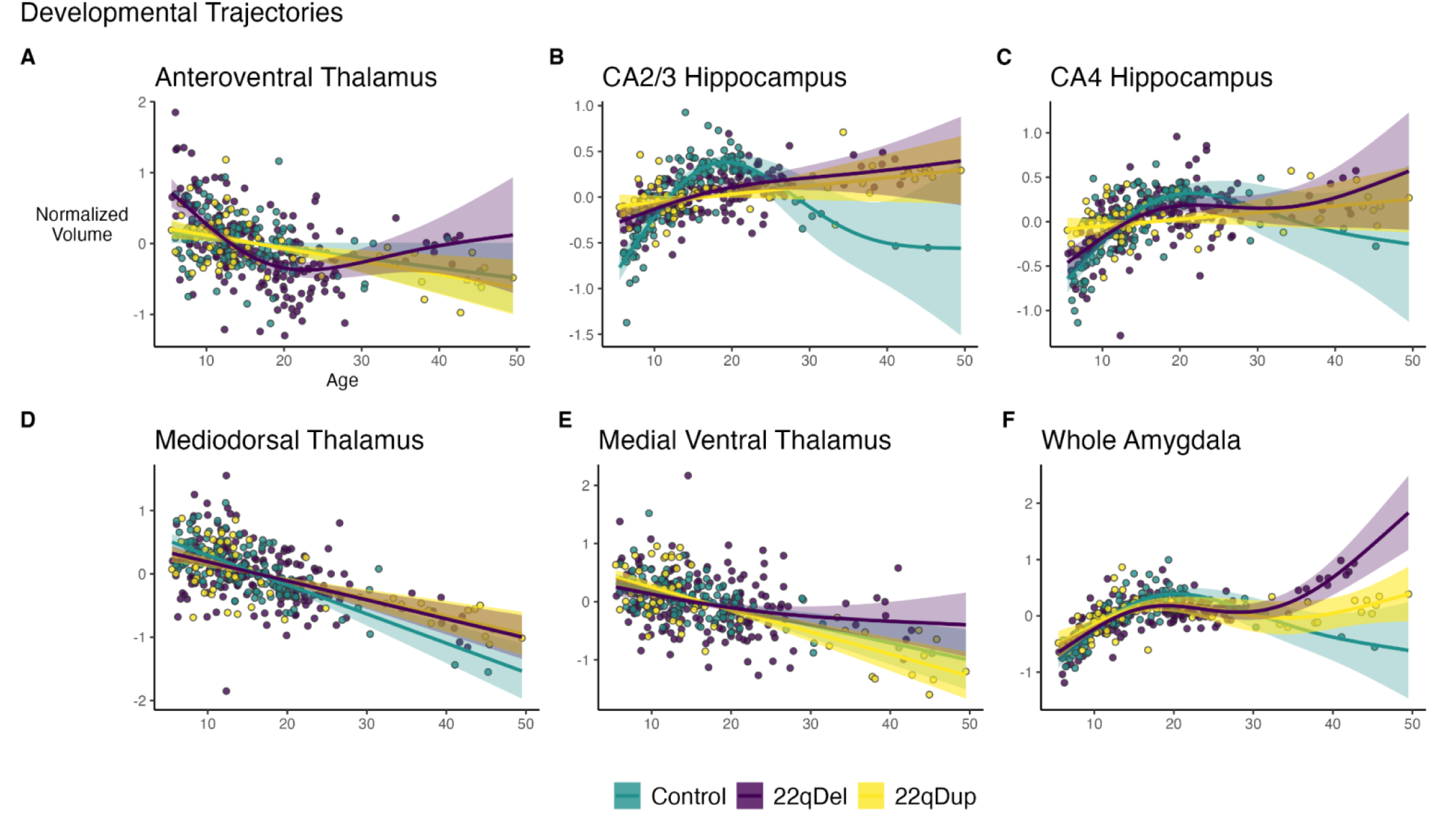
Selected age curves. GAMMs and partial residuals in 22qDel (purple), 22qDup (yellow), and TD controls (green), showing the relationship between age and selected subregion volumes, controlling for covariates. **A**) In the anteroventral thalamus, age effects are only significant in 22qDel. 22qDel volumes are significantly higher than TD between ages 5.5 and 8.7, and lower between ages 14.7 and 22.9. **B-C**) In the hippocampal CA2/3 and CA4 regions, TD controls exhibit an inverted-U-shaped developmental curve, which is mostly flattened in both 22qDel and 22qDup. CA2/3 volumes in 22qDel were greater than TD between ages 5.5 and 8.3 and lower than TD between ages 13.4 and 21.2, with similar periods of difference for 22qDup versus TD (see **Supplemental Table S4**). **D**) All groups show significant age-related decreases in mediodorsal thalamus volumes. **E**) Medial ventral (reuniens) thalamus shows similar developmental trajectories to mediodorsal thalamus, but only 22qDup had a significant main effect of age. **F**) Whole amygdala curves are similar across groups through childhood and adolescence, followed by potential increases in adulthood specific to 22qDel. Similar effects are observed for amygdala subregions.

### Secondary analyses

When tested in separate case-control analyses, 16 regions showed significant effects of 22qDel versus TD (**Table S1**), and one region (mediodorsal thalamus) showed a significant effect of 22qDup versus TD (**Table S2**), at a threshold of *q*<0.05 across the 86 tests.

Secondary analyses of separate left and right hemispheres showed strong concordance in gene dosage effects on regional volumes across both hemispheres (**Table S3**).

Significant main effects of sex on subcortical volumes were found for 28 regions (see **Table S5**). However, no significant interactions were found with gene dosage.

Hippocampal tail volume in 22qDel was significantly associated with Verbal but not Nonverbal IQ; however, no relationships were observed for 22qDup or TD groups (**Figure S3**).

No subcortical regions were found to exhibit significant relationships between volume and either continuous or categorical measures of positive psychosis-risk symptoms in the 22qDel group.

In the 22qDel group, no subcortical regions exhibited a significant relationship between antipsychotic medication status and volume. Additionally, when the gene dosage analysis was repeated with the addition of a covariate for antipsychotic medication status, the only notable change was the additional finding of a significant positive gene dosage effect in the medial geniculate nucleus of the thalamus (**Table S6**).

## Discussion

This is the first study to systematically characterize the relationship between the dosage of genomic material at the 22q11.2 locus and the volumes of specific subcortical nuclei. It is also the first study to investigate longitudinal subcortical development in 22qDup. We used an accelerated longitudinal design to recruit an unprecedented sample of 22qDel and 22qDup carriers and TD controls, spanning from childhood to middle adulthood. Using linear and nonlinear mixed effects regression approaches to map gene dosage and age effects on regional volumes, we identified several novel findings, specifically: *1)* gene dosage at the 22q11.2 locus is positively related to total intracranial and hippocampal volume, but not whole thalamus or amygdala volume; *2)* 22q11.2 gene dosage has positive relationships to specific amygdala subregions, and bi- directional relationships to specific thalamic nuclei; and, *3)* longitudinal development of subcortical structures is differentially altered in 22qDel and 22qDup across subcortical regions.

### Gene dosage effects

Standardized beta effect sizes for gene dosage effects on volume were of similar magnitude to effect sizes from previous large studies of cortical and subcortical brain structure in neurodevelopmental CNVs including 22qDel [3,19] and idiopathic neurodevelopmental and psychiatric disorders [23,24]. For example, hippocampal volume was found to be -0.46 standard deviations in patients with schizophrenia compared to controls [24], compared to our whole hippocampus effect size of -0.8 standard deviations between 22qDel and TD (**Table S1**). Case-control effect sizes were somewhat smaller in 22qDup versus TD, with the single significant region (mediodorsal thalamus; **Table S2**), exhibiting an effect size of -0.44.

These findings extend prior work from Lin et al., who provided the first evidence for a gene dosage relationship to cortical thickness, surface area, ICV, and hippocampal volume in individuals with reciprocal 22q11.2 CNVs [18]. Anatomical subregions were not investigated in that study, but a shape analysis suggested that these structures may be non-uniformly impacted. In that analysis, hippocampal thickness was found to be greater in 22qDup relative to 22qDel in regions roughly corresponding to the subiculum and CA1, which is corroborated by our current study. However, our approach did not find any regions of decreased hippocampal volume in 22qDup relative to 22qDel, whereas Lin et al. found support for some localized thickness decreases in 22qDup in regions approximately corresponding to CA2-4. Here, we expand on and broadly replicate the previous hippocampal findings in this larger longitudinal sample and have increased sensitivity to detect effects localized to specific nuclei.

Notably, within the thalamus we found bi-directional gene dosage effects across subregions. Volumes of the mediodorsal and ventral lateral nuclei decreased with increasing 22q11.2 copy number, whereas the opposite was observed for the lateral posterior, lateral geniculate and reuniens nuclei. Contrary to our hypothesis, the bi-directional effects did not follow a sensory/executive pattern; both the mediodorsal and reuniens nuclei have strong connections to the prefrontal cortex, whereas the ventral lateral, lateral posterior and lateral geniculate nuclei are more strongly connected to motor and visual cortex [52–54]. Interestingly, the reuniens nucleus, which exhibited the strongest thalamic effects, is a key hub in a network connecting the thalamus, hippocampus, and prefrontal cortex [55–57]. In 22qDel, bi-directional disruptions have been observed in functional connectivity of the thalamus and hippocampus to regions including the prefrontal cortex [58].

Analysis of hippocampal tail volume relationships to IQ suggests that this region may be particularly related to verbal cognition in 22qDel but not 22qDup or TD. This broadly supports the recent finding of hippocampal tail volume relationships with verbal learning scores in 22qDel [27]. Our analysis was specifically motivated by this prior literature; however, this effect would not have remained significant after correction for multiple comparisons in an exploratory analysis of cognition relationships to all subregional volumes.

Exploratory analyses did not find significant psychosis-risk symptom/volume relationships. However, a large multi-site study of subcortical volumes in 22qDel did find evidence for lower thalamus, hippocampus, and amygdala volumes in 22qDel individuals with psychotic disorder, compared to those without [19]. This suggests that, within the 22qDel population decreased subcortical volumes may only be detectable in those meeting full criteria for psychotic disorder, rather than subthreshold symptomatology. The current 22qDel sample was only powered to test psychosis-risk associations (**Table 1**).

### Maturational effects

Studies of normative subcortical development often show volume increases in childhood followed by decreases later in life, and this pattern is particularly prominent in the hippocampus and amygdala [59–61]. We find that in hippocampal CA2/3, and to a lesser extent CA4, both 22qDel and 22qDup failed to exhibit the expected early life increases and adult decreases observed in TD. This was not observable in the whole hippocampus, where curves overlapped across the age range. Amygdala volumes in the three groups followed more similar developmental trajectories, except that in 22qDel volumes continued to increase in adulthood, while plateauing or decreasing in the other groups. All groups exhibited similar age-related decreases across many thalamic subregions. These trajectories were more linear compared to the hippocampus and amygdala. 22qDel had abnormally steep decreases in anteroventral thalamic volumes, a thalamic subregion implicated in spatial learning and memory [62]. The steep declines in 22qDel anteroventral thalamus volumes may reflect either an abnormal developmental mechanism, or compensatory changes related to the abnormally high volume in early childhood. A prior independent longitudinal study of 22qDel and TD using a similar thalamic parcellation found an overall pattern of age-related volume decreases resembling many of the thalamic age effects in our current study [26]. However, that study used linear models rather than nonlinear splines, and as such was not sensitive to differential rates of change across different age periods, which we observe for certain regions.

Our analyses of maturational trajectories build on recent longitudinal cortical findings of altered developmental trajectories of cortical thickness and surface area in 22q11.2 CNVs [49]. Jalbrzikowski et al. found that 22qDel, 22qDup, and TD controls all showed broad decreases in cortical thickness from childhood to adulthood, but the 22qDel group showed a protracted pattern of cortical thinning. 22qDup did not exhibit the same age-related cortical surface area decreases observed in TD and 22qDel.

### Relationship to post-mortem human and animal model findings

The approach of mapping gene dosage effects on MRI-derived volumes to histologically-defined subcortical nuclei allows for more effective comparison between our neuroimaging results and findings from post-mortem brain tissue.

The strongest negative gene dosage effects were in the mediodorsal thalamus, a major source of thalamic input to the prefrontal cortex. This region has been highly studied in post-mortem brain tissue from individuals with schizophrenia, but findings are mixed, with several reporting decreased volumes and cell counts, and others reporting no differences to controls [29]. The strong effect we observe on this structure in 22q11.2 CNVs suggests a particular disruption of thalamic-prefrontal development that may be distinct from the changes underlying most idiopathic schizophrenia cases. Studies of the hippocampus in schizophrenia have demonstrated reductions in the volume and/or neuron number in subfields including the subiculum, CA1, CA2/3, and CA4 [63], which are consistent with our findings of decreased hippocampal volumes in 22qDel patients who are at increased risk for schizophrenia, compared to 22qDup who are at lower risk [14–16].

Subregion-specific analysis also helps connect results to animal model findings, which are often reported relative to these histological regions. Mouse models of 22qDel have repeatedly shown disruption to structure, function, and development of hippocampal regions CA1, CA2 and CA3 [64–66]. GABAergic inhibitory cells are particularly implicated in these disruptions. Alterations in thalamic-cortical functional connectivity in 22qDel mice have been related to changes in the auditory thalamus (medial geniculate) mediated by microRNA processes downstream of the 22q11.2 gene *Dgcr8*, which have been corroborated in human post-mortem tissue from individuals with schizophrenia [67]. Gene dosage was positively related to right medial geniculate volume in our secondary analysis of separate hemispheres (**Table S3**), and our supplemental analysis controlling for antipsychotic medication (**Table S6**). *Dgcr8* haploinsufficiency has also been linked to decreased dendritic spine density in regions including hippocampal CA1 in a 22qDel mouse model [68].

### Strengths, limitations, and future directions

Several strengths of this study should be considered in support of its reliability. The sample size of 191 scans from 96 participants with 22qDel, and 64 scans from 37 individuals with 22qDup is large for rare genetic disorders [69]. The 22qDup neuroimaging sample is unprecedented in size for that syndrome. Our large sample of 22q11.2 CNV carriers and TD controls spans a wide age range, allowing us to test important developmental hypotheses. However, the age distributions are right-skewed, and the data spanning middle adulthood were limited.

Our unique sample with identically acquired structural images from individuals with 22qDel, 22qDup, and TD controls allows for a powerful regression analysis approach in which approximate 22q11.2 gene dosage is operationalized as an integer value. This allows us to go beyond case-control differences to test specifically for brain phenotypes that are related to the content of genomic material in these reciprocal CNVs. While each carrier had a molecularly confirmed CNV at the 22q11.2 locus, breakpoints can vary in some cases [7]. However, breakpoints at this locus are largely consistent due to the pattern of low copy repeats flanking the region [9]. Breakpoint variation is thus not expected to strongly influence results. Gene dosage also likely does not fully reflect differences in gene transcription and translation in brain tissue, which are likely more closely linked to phenotypes, but cannot be measured *in vivo* or inferred from less invasively collected tissue such as blood [70,71]. When tested in separate case-control analyses, 22qDel carriers show somewhat stronger effects on subcortical structures relative to TD than do 22qDup, which is consistent with the more severe neurobehavioral/cognitive phenotype in 22qDel [69,72,73]. The smaller 22qDup sample size may partially explain the finding of fewer statistically significant differences between 22qDup and TD compared to 22qDel and TD (**Tables S1 and S2**). While the differences between 22qDel and the other two groups may be driving some of the gene dosage findings, the results of the primary gene dosage models along with the supplemental case-control analyses are highly informative, highlighting that the overall trend for subcortical regions is towards linear gene dosage relationships to volume, rather than convergent effects in 22qDel and 22qDup. In other words, there are no regions for which 22qDel and 22qDup significantly differ from TD in the same direction.

Future directions of research include characterization of gene dosage effects in other reciprocal CNVs such as the 16p11.2 and 15q11-q13 loci and mapping subcortical development in individuals with idiopathic autism and those at clinical high risk (CHR) for psychosis to determine convergent (and/or divergent) patterns. Larger multi-site studies of 22q11.2 CNVs will be better-powered to elucidate possible roles of breakpoint variation, and brain-behavior relationships. Analysis of cytoarchitecture and gene expression in post-mortem tissue from individuals with 22q11.2 CNVs will also be highly informative. Research in animal and *in vitro* models will continue to bridge the gap to understanding circuit-level dysfunction, and the roles of individual genes and potential targeted pharmacological interventions.

## Conclusions

This study is the first to characterize gene dosage effects on subregional subcortical volumes in individuals with reciprocal 22q11.2 CNVs, and the first study of longitudinal development of subcortical structures in 22qDup. Using a linear mixed effects approach, we found positive gene dosage effects on total brain volume, hippocampal volumes, and several amygdala subregions, and bi-directional effects across multiple thalamic nuclei. GAMM analyses revealed both distinct and convergent disruptions in developmental trajectories in 22q11.2 CNVs across the thalamus, hippocampus, and amygdala. These results highlight the impact of genes in the 22q11.2 locus on subcortical brain development and motivate future research linking gene expression to brain phenotypes at the levels of cells, circuits, and macro-scale structures.

## Author Contributions

**Charles H. Schleifer**: Conceptualization, Methodology, Software, Validation, Formal Analysis, Data Curation, Writing - Original Draft, Writing - Review & Editing, Visualization

**Kathleen P. O’Hora**: Data Curation, Methodology, Writing - Review & Editing

**Hoki Fung**: Data Curation, Methodology, Writing - Review & Editing

**Jenny Xu**: Data Curation, Writing - Review & Editing

**Taylor-Ann Robinson**: Data Curation

**Angela S. Wu**: Data Curation, Writing - Review & Editing

**Leila Kushan-Wells**: Investigation, Resources, Data Curation, Project Administration

**Amy Lin**: Data Curation

**Christopher R. K. Ching**: Conceptualization, Writing - Review & Editing

**Carrie E. Bearden**: Conceptualization, Investigation, Writing - Review & Editing, Supervision, Project Administration, Funding Acquisition.

## Data availability

Data are publicly available from the National Institute of Mental Health Data Archives: https://nda.nih.gov/edit_collection.html?id=2414

To facilitate reproducibility and rigor, analysis code is publicly available on GitHub: https://github.com/charles-schleifer/22q_subcort_volumes

