## Supplement for "Effects of Gene Dosage and Development on Subcortical Nuclei Volumes in Individuals with 22q11.2 Copy Number Variations"

**Supplemental Methods**

Participants

The total longitudinal sample consisted of 387 scans from 213 participants (5.5–49.5 years of age; n=96 22qDel baseline, 53.1% female; n=37 22qDup baseline, 45.9% female; n=80 TD controls baseline, 51.3% female), recruited from an ongoing longitudinal study at the University of California, Los Angeles (UCLA). The 22qDel and 22qDup participants all had a molecularly confirmed 22q11.2 CNV. Participants had data from between 1 and 6 time points (mean=1.81 visits, SD=1.04), separated by an average of approximately one- and three-quarter years (mean=1.76 years, SD=1.16). The three groups were statistically matched based on baseline age and sex, as well as mean number of longitudinal visits and interval between visits, using appropriate tests (ANOVA, or chi-squared). Exclusion criteria for all study participants were as follows: significant neurological or medical conditions (unrelated to 22q11.2 deletion or duplication) that might affect brain structure, history of head injury with loss of consciousness, insufficient fluency in English, and/or substance or alcohol use disorder within the past 6 months. As we aimed to include a representative cohort of CNV carriers, patients with cardiac-related and/or immune issues were not excluded, as these are common medical comorbidities in 22qDel. Healthy controls were free from significant intellectual disability and/or family history of psychotic disorder, and did not meet criteria for any psychiatric disorder, with the exception of attention deficit-hyperactivity disorder, anxiety disorders, or a past episode of depression, due to their prevalence in childhood and adolescence [1–3]. After study procedures had been fully explained, adult participants provided written consent, while participants under the age of 18 years provided written assent with the written consent of their parent or guardian. The UCLA Institutional Review Board approved all study procedures and informed consent documents.

*Clinical assessment*

At each study time point, demographic information and clinical measures were collected for each participant by trained Master's-level clinicians, supervised by a licensed clinical psychologist. Psychiatric diagnoses were established with the Structured Clinical Interview for DSM-IV (SCID), with an additional developmental disorders module [4]. Verbal IQ was assessed via the Wechsler Abbreviated Scale of Intelligence (WASI-2) Vocabulary subtest, and nonverbal IQ was assessed via the WASI-2 Matrix Reasoning subtest. Dimensional psychosis-risk and general psychiatric symptoms were assessed via the Structured Interview for Psychosis-Risk Syndromes (SIPS) [5]. For more details on study ascertainment and recruitment procedures, see Jalbrzikowski et al. 2012 and 2013 [6,7].

*Neuroimaging acquisition*

All subjects were imaged at the UCLA Center for Cognitive Neuroscience on either a Siemens TimTrio scanner (with a 12-channel head coil) or Siemens Prisma (with a 32-channel head coil). T1w scans were acquired in sagittal slices with 1mm$^3$ voxels, as described in Jalbrzikowski et al. 2022 [8], using MPRAGE sequences adapted from the Alzheimer's Disease Neuroimaging Initiative (ADNI) protocol [9]. Trio and Prisma MPRAGE scans used nearly identical parameters: TR = 2.3 s, FOV = 256 mm, matrix = 240 × 256, flip angle = 9°, slice thickness = 1.20 mm, 160 slices. TE was 2.91 ms for Trio scans, and 2.94 ms for Prisma.

*Neuroimaging preprocessing*

T1w MRI scans were processed with the FreeSurfer analysis package, version 7.3.2 [10]. Scan sessions at all timepoints were first processed cross-sectionally using the recon-all anatomical segmentation pipeline [11,12]. The FreeSurfer longitudinal stream was subsequently applied, which has been shown to significantly improve reliability and statistical power in repeated-measure analyses [13]. This method generates unbiased within-subject templates using robust, inverse consistent registration, and uses these templates to improve initialization of several processing steps, such as skull-stripping, Talairach transforms, atlas registration, spherical surface maps, and parcellations [13].

*Subcortical nuclei volumes*

For each MRI scan, the FreeSurfer longitudinal segment subregions pipeline was used to estimate volumes for 25 thalamic subregions, 19 hippocampal subregions, and 9 amygdala subregions, as well as whole-structure volumes [14–17]. These methods use Bayesian inference to automatically segment T1w MRI images using probabilistic template atlases based on histological data and ultra-high-resolution *ex vivo* MRI. These segmentations have been highly validated and have been applied by multiple groups and consortia to large scale neuroimaging analyses of development and group differences in various psychiatric conditions [18–22]. For analysis, several hippocampal subregions were combined as follows: the head and tail of hippocampal CA1 were added to give a single CA1 volume, similarly for CA2/3, CA4, molecular layer, GC-ML-DG, presubiculum, and subiculum, as in Mancini et al. 2010, and Latrèche et al. 2023) [23,24]. Hippocampal regions CA2 and CA3 are combined into a single CA2/3 region in the FreeSurfer segment subregions atlas due to difficulty reliably determining the boundary between regions [14]. In the thalamus several regions were combined, as in Huang et al. 2020 [21]: the mediodorsal medial and lateral regions were combined to give one mediodorsal region; the ventral lateral anterior and posterior subregions were combined into one ventral lateral region; the ventral anterior and ventral anterior magnocellular regions were combined into one ventral anterior region; the anterior, lateral, and inferior pulvinar were combined into one pulvinar region. For the primary analyses, regional volumes were averaged within subjects between left and right hemispheres to reduce multiple comparisons and facilitate interpretation. See **Supplemental Results Table S1** for bilateral gene dosage analyses without this averaging step.

An example image was generated for visualization purposes (see **Figure S2**) using the FreeSurfer recon-all and segment subregions pipelines with the MNI152 template brain as the input [25].

*Quality control*

Several qualitative and quantitative approaches were taken to prevent inclusion of inaccurately estimated volumes in the analysis. First, each raw T1w image was visually assessed for quality prior to preprocessing, and excluded if quality was low (e.g., significant motion artifact, signal loss, or incomplete brain coverage). After subcortical segmentation, each image was visually checked to ensure alignment of thalamus, hippocampus, and amygdala masks with their associated structures. In four scans, bilateral thalamic volumes were excluded from further analysis because the thalamic segmentation was found to include parts of the striatum. We also excluded the medial pulvinar from further analysis in all subjects because the boundaries of the medial pulvinar mask were observed to extend beyond the thalamus in many cases. Finally, we calculated the mean and standard deviation of each subregion volume in the full cohort, and for each individual, excluded a given subregion from further analysis if the volume was greater than 3 standard deviations absolute difference from the overall group mean. Out of the total set of 33,196 regions from 386 subjects, 139 total regions across 67 subjects were flagged for exclusion by this metric. For the main analyses, volumes were averaged bilaterally except in the cases where a region had been excluded as an outlier in one hemisphere, in which case the non-outlier volume was used.

*Data harmonization*

To harmonize data acquired on two different scanners, we applied a longitudinal implementation of the ComBat algorithm using the longComBat package in R version 4.2.2 [26,27]. ComBat uses empirical Bayes methods to estimate and remove scanner/batch effects with increased robustness to outliers in small samples compared to general linear model approaches. ComBat was initially developed for genomics data [27], and has been subsequently adapted for neuroimaging and shown to preserve biological associations while effectively removing unwanted non-biological variation associated with site/scanner [28]. The longitudinal adaptation, which uses random effects to account for within-subject repeated measures, has been shown to further increase statistical power in longitudinal neuroimaging analyses [26]. LongComBat has been used to harmonize structural and functional MRI features in largely overlapping cohorts of individuals with 22qDel and controls [8,29]. LongComBat requires that the input data matrix not contain missing values, so for regions that were to be excluded from the final analysis, we imputed values based on the mean of that region’s volume in individuals collected on the same scanner (Trio or Prisma) with the same CNV status (22qDel, 22qDup, or TD). Excluded volumes were then set to “NA” after longComBat harmonization, prior to regression analysis.

*Gene dosage effects on volume*

To investigate the impact of CNV status on subcortical volumes, we used a linear mixed effects approach where gene dosage at the 22q11.2 locus was numerically coded based on CNV status: 22qDel=1, TD=2, and 22qDup=3 copies of the locus. First, total intracranial volume (ICV) was normalized based on the TD group mean and standard deviation, and a linear random effects model was tested predicting total ICV from gene dosage, controlling for linear and quadratic age, sex, and scanner, with a random intercept for subject ID to account for repeat visits. Models were fit with restricted estimation of maximum likelihood (REML) and *p*-values were computed with the Kenward-Roger approximation for degrees of freedom [30] using the lmerTest package in R [31]. Gene dosage effects were first tested for the whole thalamus, hippocampus, and amygdala. Then, we tested each of the 40 subregions, computed p-values for the effect of gene dosage, and corrected for multiple comparisons with False Discovery Rate (FDR) [32].

*Age effects*

To characterize developmental trajectories of subcortical volumes, we assessed nonlinear age-related changes in each cohort with general additive mixed models (GAMMs) as in Jalbrzikowski et al., 2022 [8] using the mgcv package in R [33]. GAMMs are a nonlinear extension of mixed effects models, and as such are well-suited to data with repeated measures within subjects. Regression splines are used to estimate nonlinear curves, and overfitting is prevented through REML and penalization of higher order polynomials [33–35]. An advantage of this approach is that curves can take different forms in each cohort. For each normalized subcortical volume, a GAMM was computed predicting volume from age in each group, controlling for sex, total brain volume, and scanner, with a random intercept for subject ID. p-values for the effect of age in each group were FDR-corrected for multiple comparisons. Age ranges of significant difference between CNV carrier groups and TD controls were computed based on the 95% confidence interval for the difference in smoothed curves.

*Secondary analyses*

Several secondary analyses were performed to complement the primary gene dosage analyses. Regional volume differences compared to the TD group were tested separately for 22qDel and 22qDup groups. Gene dosage analyses were repeated with antipsychotic medication status as an additional covariate. Additional models of antipsychotic status effects on volume were tested in only 22qDel, which was the only group with multiple more than 10% of participants taking antipsychotic medication. Gene dosage analyses were also repeated without averaging structures bilaterally to detect any asymmetric hemispheric effects. Secondary analyses of the interaction between sex and gene dosage on brain volumes were also tested.

Motivated by existing literature relating low hippocampal tail volume to verbal learning impairment in 22qDel [23], we assessed verbal and non-verbal IQ (Wechsler Abbreviated Scale of Intelligence (WASI-2) Vocabulary and Matrix subtest scaled scores) for associations with hippocampal tail volume in each group. A recent study of hippocampal volumes in 22qDel from another research group found that decreased hippocampal tail volume was associated with impaired development of verbal learning [23]. Here we sought to replicate that finding and extend to 22qDup. In each group (22qDel, 22qDup, and TD) we tested a linear mixed model predicting verbal and non-verbal IQ (WASI-2 Vocabulary Verbal and Matrix Reasoning subtest scaled scores) from hippocampal tail volume, controlling for sex and scanner, with a random intercept for subject ID. See **Supplemental Figure S3** for results.

Because 22qDel is associated with psychosis risk, we tested relationships between psychosis risk symptoms and subcortical volumes in the 22qDel group. Specifically, we tested models relating positive symptom scores from the Structured Interview for Psychosis-Risk Syndromes (SIPS) [5] to volume across each region, controlling for age, $age^2$, sex, and scanner, with a random intercept for each individual participant. We also similarly tested models relating volume to categorical diagnosis of Psychosis Risk Symptoms, operationalized here as having any score of 3 or greater (i.e., prodromal range) on any SIPS positive symptom item.

### Supplemental Results

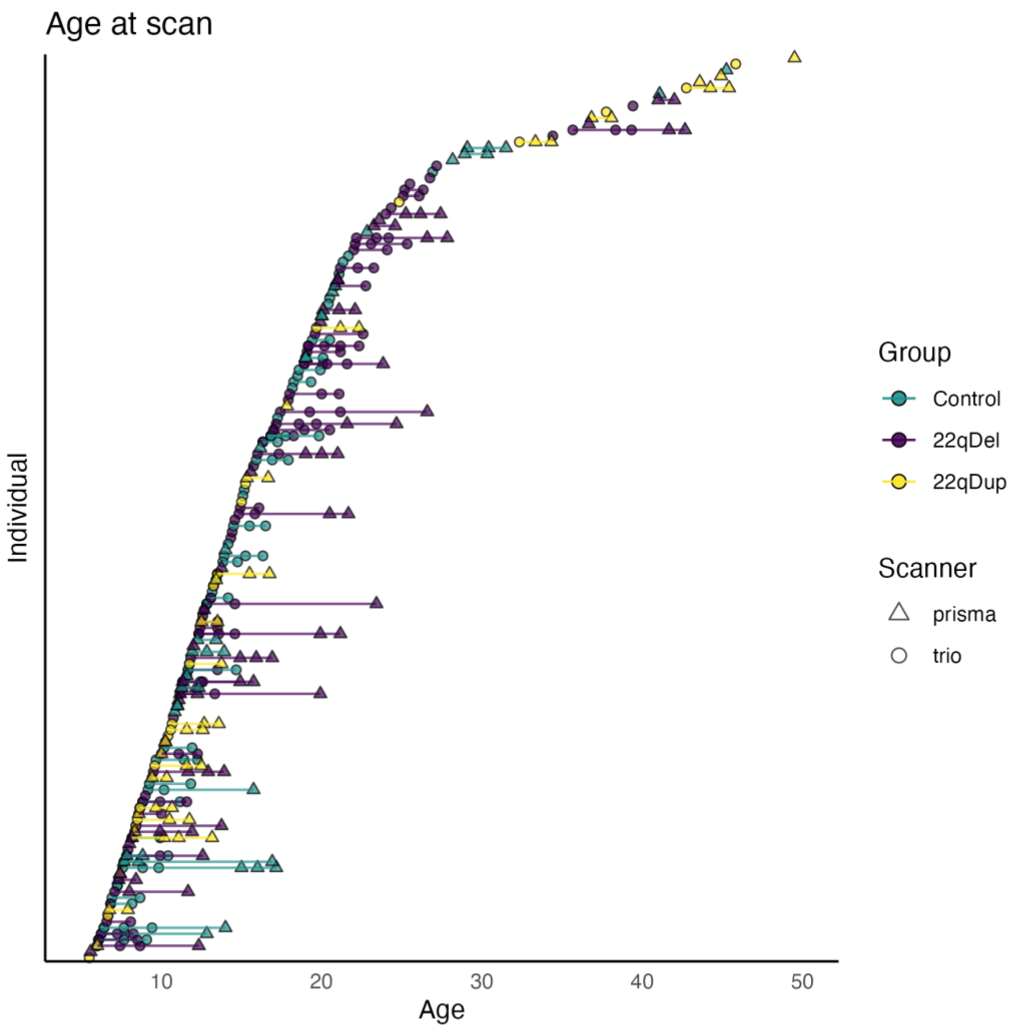


**Figure S1. Participant age distribution.** Typically developing controls in green, 22qDel in purple, and 22qDup in yellow, with lines connecting follow-up visits from the same individual. Scanner type (Siemens Trio or Prisma) indicated by circle or triangle, respectively.

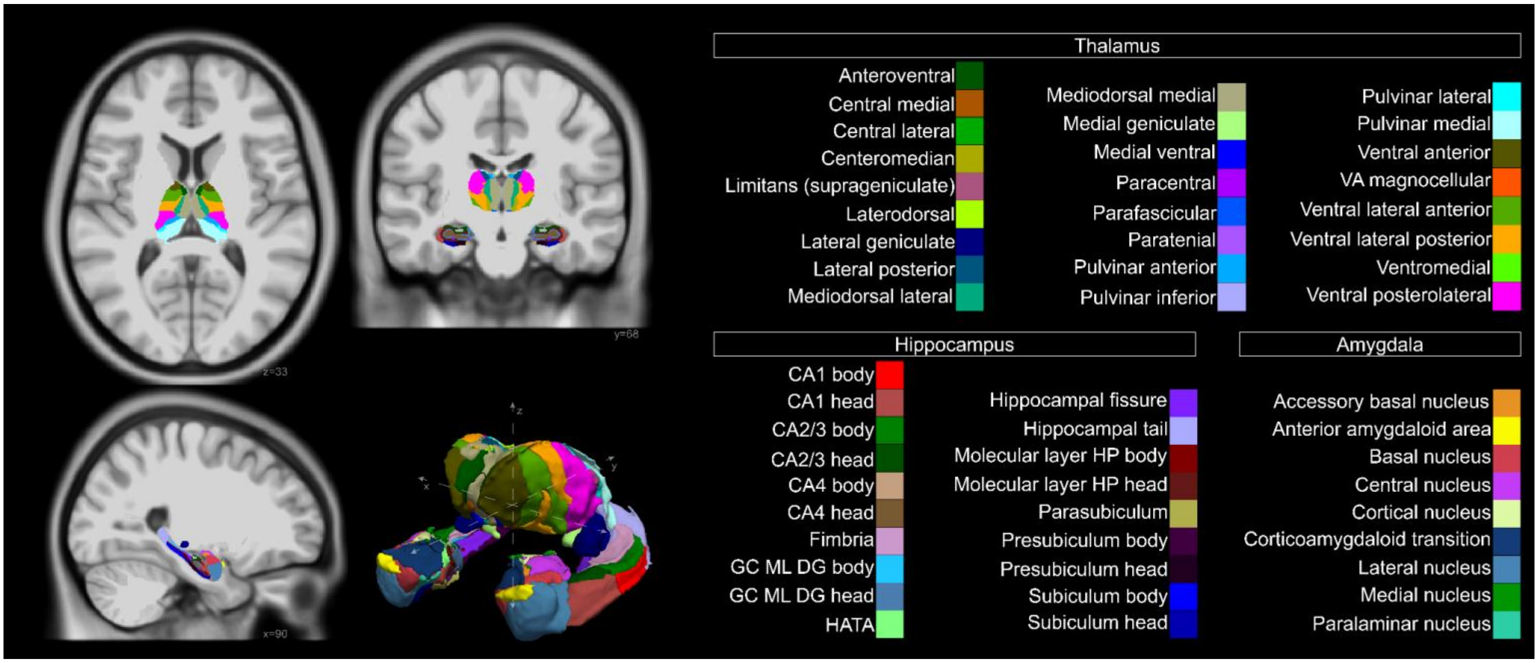


**Figure S2**. **Anatomical parcellation of thalamus, hippocampus, and amygdala nuclei.** *Left:* axial, coronal, and sagittal slices, and 3D reconstruction of structures. Generated for visualization purposes using the FreeSurfer recon-all and segment subregions pipelines with the MNI152 template brain as the input. *Right:* structure names and color key. Abbreviations: VA = Ventral Anterior, CA = Cornu Ammonis (areas 1, 3 and 4), GC ML DG = Granule Cell and Molecular Layer of the Dentate Gyrus, HATA = Hippocampus Amygdala Transition Area, HP = Hippocampus.

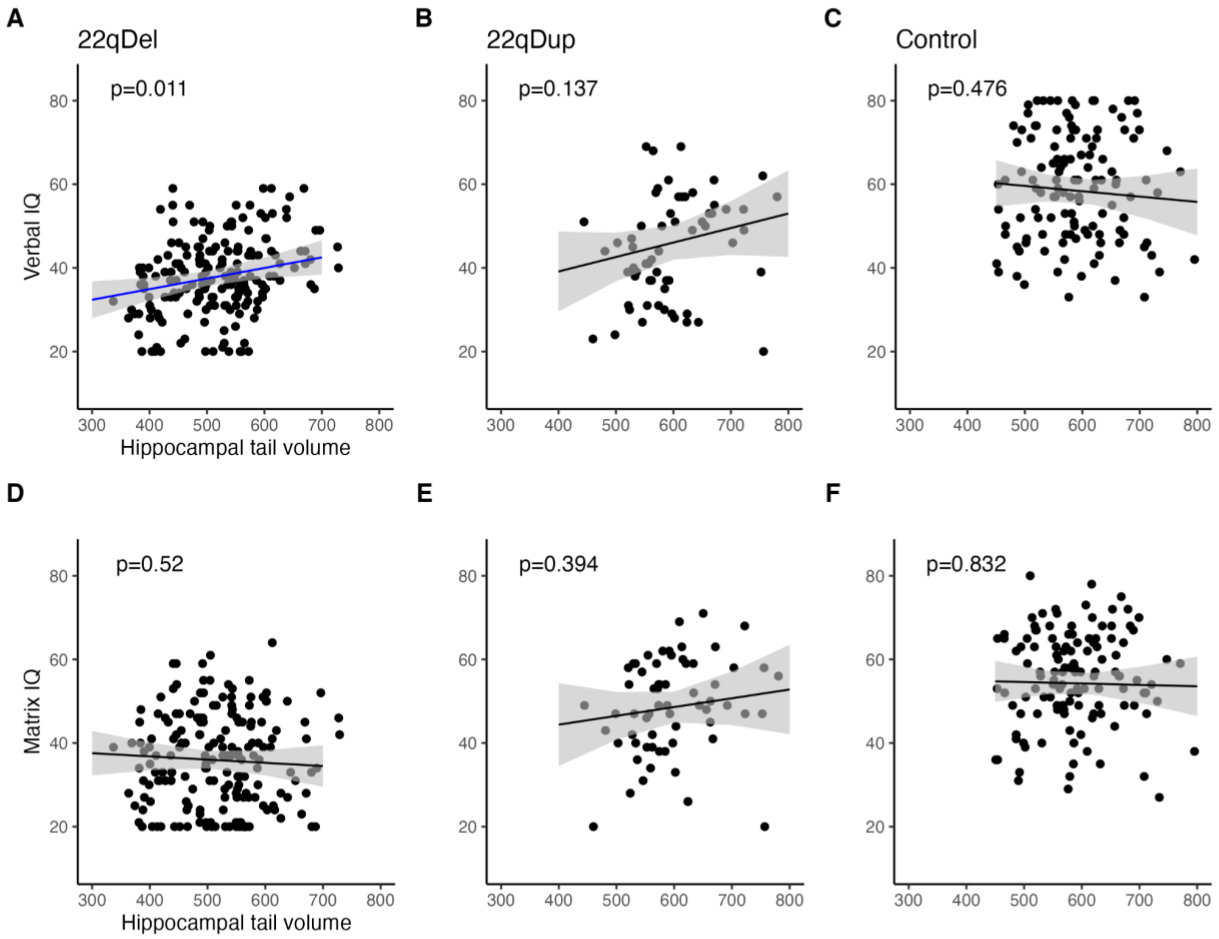


**Figure S3. Relationships between IQ subtests and hippocampal tail volume. A-C**) WASI Verbal IQ scaled scores were significantly predicted by hippocampal tail volume controlling for sex, site, and participant in 22qDel (*beta*=1.86, *p*=0.011) but not 22qDup or TD controls. **D-F**) Matrix Reasoning (Nonverbal IQ) subscale scores were not related to hippocampal tail volume in any group. Adding Nonverbal IQ as a covariate in the model predicting Verbal IQ from hippocampal tail volumes increased the strength of the verbal IQ relationship in 22qDel (*beta*=2.07, *p*=0.0018).

| Structure | Region | *beta* | *p* | FDR *q* |
|---|---|---|---|---|
| thalamus | lateral geniculate | -0.74 | 3.60E-07 | 6.20E-06 |
| | medial ventral (reuniens) | -0.61 | 7.50E-05 | 0.00065 |
| | medial geniculate | -0.32 | 0.0053 | 0.033 |
| | parafascicular | -0.32 | 0.0067 | 0.036 |
| | ventral lateral | 0.28 | 0.018 | 0.081 |
| | mediodorsal | 0.29 | 0.031 | 0.12 |
| | lateral posterior | -0.23 | 0.11 | 0.33 |
| | central medial | -0.23 | 0.14 | 0.39 |
| | paracentral | 0.2 | 0.18 | 0.4 |
| | limitans (suprageniculate) | 0.18 | 0.19 | 0.42 |
| | anteroventral | -0.18 | 0.19 | 0.42 |
| | laterodorsal | 0.14 | 0.31 | 0.57 |
| | ventral anterior | -0.07 | 0.53 | 0.78 |
| | whole thalamus | -0.05 | 0.67 | 0.88 |
| | pulvinar | -0.05 | 0.71 | 0.88 |
| | paratenial | -0.05 | 0.73 | 0.88 |
| | ventral posterolateral | -0.04 | 0.74 | 0.88 |
| | ventromedial | -0.02 | 0.85 | 0.94 |
| | centromedian | -0.02 | 0.87 | 0.94 |
| | central lateral | -0.01 | 0.92 | 0.96 |
| hippocampus | subiculum | -0.91 | 3.60E-11 | 3.10E-09 |
| | hippocampal tail | -0.99 | 1.80E-10 | 7.80E-09 |
| | molecular layer | -0.87 | 2.40E-08 | 6.80E-07 |
| | whole hippocampus | -0.8 | 4.10E-08 | 8.80E-07 |
| | GC ML DG | -0.73 | 7.80E-07 | 1.10E-05 |
| | CA4 | -0.7 | 1.80E-06 | 2.20E-05 |
| | CA1 | -0.73 | 4.50E-06 | 4.90E-05 |
| | hippocampal fissure | -0.58 | 6.20E-05 | 0.00059 |
| | presubiculum | -0.22 | 0.13 | 0.36 |
| | parasubiculum | 0.23 | 0.15 | 0.39 |
| | CA2/3 | -0.2 | 0.16 | 0.4 |
| | hippocampal amygdala transition area | 0.2 | 0.18 | 0.4 |
| | fimbria | -0.14 | 0.35 | 0.59 |
| amygdala | basal nucleus | -0.52 | 0.00016 | 0.0012 |
| | paralaminar nucleus | -0.48 | 0.00051 | 0.0037 |
| | medial nucleus | -0.46 | 0.00098 | 0.0065 |
| | accessory basal nucleus | -0.38 | 0.0064 | 0.036 |
| | whole amygdala | -0.33 | 0.012 | 0.059 |
| | cortical nucleus | -0.33 | 0.021 | 0.089 |
| | central nucleus | -0.31 | 0.034 | 0.13 |
| | lateral nucleus | -0.15 | 0.26 | 0.54 |
| | anterior amygdaloid area | -0.12 | 0.34 | 0.59 |
| | corticoamygdaloid transition | 0.12 | 0.39 | 0.62 |

**Table S1. 22qDel versus TD comparisons.** Linear mixed models predicting normalized volume from group (22qDel or TD), controlling for age, age $^2$, total brain volume, sex, scanner, and subject ID. False Discovery Rate (FDR) correction was applied across all 86 tests for 22qDel and 22qDup comparisons to TD.

| Structure | Region | *beta* | *p* | FDR *q* |
|---|---|---|---|---|
| thalamus | mediodorsal | -0.44 | 0.0079 | 0.04 |
| | ventral posterolateral | -0.29 | 0.1 | 0.33 |
| | limitans (suprageniculate) | 0.3 | 0.12 | 0.35 |
| | laterodorsal | 0.29 | 0.16 | 0.4 |
| | pulvinar | 0.27 | 0.18 | 0.4 |
| | lateral posterior | 0.22 | 0.26 | 0.54 |
| | ventromedial | -0.2 | 0.27 | 0.54 |
| | ventral lateral | -0.17 | 0.28 | 0.54 |
| | paracentral | -0.19 | 0.29 | 0.55 |
| | central lateral | 0.22 | 0.32 | 0.59 |
| | lateral geniculate | -0.15 | 0.4 | 0.63 |
| | paratenial | -0.14 | 0.44 | 0.67 |
| | medial geniculate | -0.08 | 0.61 | 0.85 |
| | whole thalamus | -0.06 | 0.68 | 0.88 |
| | medial ventral (reuniens) | 0.07 | 0.71 | 0.88 |
| | anteroventral | 0.07 | 0.71 | 0.88 |
| | ventral anterior | -0.05 | 0.74 | 0.88 |
| | central medial | 0.04 | 0.85 | 0.94 |
| | centromedian | 0.03 | 0.88 | 0.94 |
| | parafascicular | 0 | 0.98 | 0.98 |
| hippocampus | hippocampal fissure | 0.46 | 0.022 | 0.089 |
| | presubiculum | -0.35 | 0.07 | 0.25 |
| | hippocampal amygdala transition area | 0.25 | 0.16 | 0.4 |
| | CA4 | 0.17 | 0.34 | 0.59 |
| | CA2/3 | 0.18 | 0.36 | 0.59 |
| | fimbria | 0.19 | 0.36 | 0.59 |
| | CA1 | 0.16 | 0.42 | 0.64 |
| | subiculum | -0.12 | 0.51 | 0.76 |
| | GC ML DG | 0.09 | 0.61 | 0.85 |
| | whole hippocampus | 0.05 | 0.76 | 0.9 |
| | parasubiculum | 0.05 | 0.8 | 0.94 |
| | molecular layer | 0.01 | 0.94 | 0.96 |
| | hippocampal tail | 0.01 | 0.98 | 0.98 |
| amygdala | cortical nucleus | -0.34 | 0.077 | 0.26 |
| | medial nucleus | -0.36 | 0.084 | 0.28 |
| | anterior amygdaloid area | -0.16 | 0.34 | 0.59 |
| | corticoamygdaloid transition | 0.11 | 0.55 | 0.78 |
| | accessory basal nucleus | 0.08 | 0.65 | 0.87 |
| | basal nucleus | 0.08 | 0.65 | 0.87 |
| | central nucleus | -0.05 | 0.82 | 0.94 |
| | whole amygdala | 0.03 | 0.86 | 0.94 |
| | lateral nucleus | 0.02 | 0.91 | 0.96 |
| | paralaminar nucleus | -0.02 | 0.91 | 0.96 |

**Table S2. 22qDup versus TD comparisons.** Linear mixed models predicting normalized volume from group (22qDup or TD), controlling for age, age $^2$, total brain volume, sex, site, and subject ID. False Discovery Rate (FDR) correction was applied across all 86 tests for 22qDel and 22qDup comparisons to TD.

| Structure | Region | Hemi | *beta* | *p* | FDR *q* |
|---|---|---|---|---|---|
| whole volumes | total ICV | NA | 0.32 | 0.00012 | 0.00025 |
| | whole thalamus | L | -0.08 | 0.28 | 0.29 |
| | whole thalamus | R | 0.01 | 0.88 | 0.88 |
| | whole amygdala | L | 0.14 | 0.1 | 0.11 |
| | whole amygdala | R | 0.14 | 0.072 | 0.079 |
| | whole hippocampus | L | 0.43 | 3.70E-06 | 1.20E-05 |
| | whole hippocampus | R | 0.47 | 2.70E-07 | 2.00E-06 |
| thalamus subregions | mediodorsal | L | -0.34 | 4.50E-05 | 0.00011 |
| | mediodorsal | R | -0.32 | 0.00017 | 0.00031 |
| | ventral lateral | R | -0.28 | 0.00017 | 0.00031 |
| | ventral lateral | L | -0.26 | 0.00056 | 0.00092 |
| | paracentral | R | -0.2 | 0.017 | 0.02 |
| | medial geniculate | R | 0.17 | 0.017 | 0.02 |
| | lateral geniculate | L | 0.21 | 0.019 | 0.022 |
| | anteroventral | L | 0.24 | 0.0072 | 0.0094 |
| | medial ventral (reuniens) | R | 0.34 | 0.00064 | 0.001 |
| | medial ventral (reuniens) | L | 0.42 | 6.90E-05 | 0.00016 |
| | lateral geniculate | R | 0.46 | 9.80E-07 | 5.30E-06 |
| hippocampus subregions | fimbria | R | 0.27 | 0.0055 | 0.0077 |
| | GC ML DG | L | 0.37 | 0.00013 | 0.00025 |
| | CA1 | L | 0.38 | 0.00028 | 0.00049 |
| | CA4 | L | 0.39 | 6.30E-05 | 0.00015 |
| | hippocampal fissure | R | 0.39 | 3.30E-05 | 8.90E-05 |
| | CA4 | R | 0.43 | 1.80E-06 | 7.60E-06 |
| | GC ML DG | R | 0.44 | 2.20E-06 | 8.40E-06 |
| | subiculum | R | 0.45 | 3.20E-07 | 2.00E-06 |
| | molecular layer | L | 0.46 | 2.90E-06 | 1.00E-05 |
| | subiculum | L | 0.47 | 1.40E-07 | 1.30E-06 |
| | molecular layer | R | 0.48 | 1.40E-06 | 6.70E-06 |
| | CA1 | R | 0.49 | 4.00E-06 | 1.20E-05 |
| | hippocampal tail | R | 0.55 | 5.00E-08 | 6.30E-07 |
| | hippocampal fissure | L | 0.57 | 7.10E-10 | 1.40E-08 |
| | hippocampal tail | L | 0.61 | 2.20E-10 | 8.20E-09 |
| amygdala subregions | paralaminar nucleus | R | 0.23 | 0.012 | 0.015 |
| | accessory basal nucleus | L | 0.23 | 0.019 | 0.022 |
| | paralaminar nucleus | L | 0.25 | 0.0059 | 0.008 |
| | basal nucleus | R | 0.27 | 0.00077 | 0.0012 |
| | basal nucleus | L | 0.29 | 0.0016 | 0.0024 |

**Table S3. Gene dosage effects in individual hemispheres.** Repeat of main analyses without averaging regions bilaterally, showing highly similar effects of gene dosage on regional volume in the left and right hemispheres. Regions with FDR $q < 0.05$ are listed in this table.

| Structure | Region | diff_TD_22qDel | diff_TD_22qDup |
| --- | --- | --- | --- |
| thalamus | anteroventral | 5.5-8.7\|14.7-22.9 | |
| | laterodorsal | | |
| | medial geniculate | | |
| | medial ventral (reuniens) | | |
| | mediodorsal | | |
| | parafascicular | 14.4-21\|39.6-49.5 | 5.5-9.9\|15.3-24.4 |
| | ventromedial | 14.9-20.4 | 5.5-11.1\|15.8-26 |
| hippocampus | CA1 | | |
| | CA2/3 | 5.5-8.3\|13.4-21.2 | 5.5-9.3\|13.4-22.2 |
| | CA4 | | 5.5-9.3\|14.8-22.5 |
| | GC ML DG | | 5.5-8.8\|14.6-22.2 |
| | hippocampal amygdala transition area | | |
| | hippocampal tail | | |
| | molecular layer | | 14.6-18.8 |
| | subiculum | | 16.6-20 |
| | whole hippocampus | | |
| amygdala | anterior amygdaloid area | 35.5-49.5 | |
| | basal nucleus | 19.1-19.6 | |
| | central nucleus | 10-19\|24-46.6 | 12-19.8\|28.4-40.4 |
| | corticoamygdaloid transition | 15.4-22.1 | 5.5-8.2\|15.6-23.1 |
| | lateral nucleus | 18.7-26.3\|42.9-49.5 | 5.5-10.3\|14.2-23.6 |
| | whole amygdala | 17.4-21\|39-49.5 | |

**Table S4. Age ranges with significant differences between CNV carriers and controls.** Age periods with group difference based on 95% confidence interval (CI), multiple discontinuous ranges separated with "|". diff_TD_22qDel lists differences between 22qDel and TD curves, diff_TD_22qDup shows the same for 22qDup.

| Structure | Region | *beta* | *p* | FDR *q* |
|---|---|---|---|---|
| thalamus | mediodorsal | 0.33 | 1.20E-02 | 0.021 |
| | anteroventral | 0.34 | 1.80E-02 | 0.028 |
| | ventromedial | 0.34 | 6.10E-03 | 0.012 |
| | ventral anterior | 0.37 | 1.60E-03 | 0.0038 |
| | paratenial | 0.37 | 6.30E-03 | 0.012 |
| | parafascicular | 0.4 | 7.20E-04 | 0.0018 |
| | ventral lateral | 0.4 | 6.80E-04 | 0.0018 |
| | centromedian | 0.42 | 6.90E-04 | 0.0018 |
| | whole thalamus | 0.55 | 8.90E-06 | 0.00019 |
| | pulvinar | 0.6 | 9.00E-05 | 0.00048 |
| hippocampus | CA2/3 | 0.31 | 2.50E-02 | 0.038 |
| | subiculum | 0.38 | 3.60E-03 | 0.0074 |
| | parasubiculum | 0.39 | 1.50E-02 | 0.024 |
| | fimbria | 0.45 | 2.30E-03 | 0.0049 |
| | HATA | 0.49 | 6.70E-04 | 0.0018 |
| | CA4 | 0.5 | 3.30E-04 | 0.0014 |
| | molecular layer | 0.5 | 5.40E-04 | 0.0018 |
| | GC ML DG | 0.52 | 1.80E-04 | 0.00088 |
| | CA1 | 0.53 | 5.20E-04 | 0.0018 |
| | whole hippocampus | 0.54 | 7.20E-05 | 0.00044 |
| | presubiculum | 0.6 | 2.80E-05 | 0.00024 |
| amygdala | anterior amygdaloid area | 0.31 | 1.40E-02 | 0.024 |
| | accessory basal nucleus | 0.43 | 1.80E-03 | 0.0041 |
| | corticoamygdaloid transition | 0.48 | 4.50E-04 | 0.0017 |
| | basal nucleus | 0.54 | 3.70E-05 | 0.00026 |
| | whole amygdala | 0.56 | 1.50E-05 | 0.00021 |
| | lateral nucleus | 0.56 | 2.00E-05 | 0.00021 |
| | paralaminar nucleus | 0.69 | 6.60E-07 | 2.80E-05 |

**Table S5. Main effects of sex**. Regions with a FDR significant main effect of sex in the model predicting volume from gene dosage, sex, age, age$^2$, scanner, and subject ID.

| Structure | Region | *beta* | *p* | FDR *q* |
|---|---|---|---|---|
| thalamus | mediodorsal | -0.33 | 6.80E-05 | 0.00029 |
| | ventral lateral | -0.26 | 5.00E-04 | 0.0016 |
| | medial geniculate | 0.18 | 1.40E-02 | 0.039 |
| | lateral posterior | 0.22 | 1.80E-02 | 0.045 |
| | lateral geniculate | 0.36 | 6.50E-05 | 0.00029 |
| | medial ventral (reuniens) | 0.39 | 7.60E-05 | 0.0003 |
| hippocampus | GC ML DG | 0.43 | 2.70E-06 | 1.50E-05 |
| | CA4 | 0.44 | 1.60E-06 | 1.10E-05 |
| | CA1 | 0.47 | 2.10E-06 | 1.30E-05 |
| | whole hippocampus | 0.48 | 1.20E-07 | 1.30E-06 |
| | subiculum | 0.5 | 8.70E-09 | 1.20E-07 |
| | molecular layer | 0.51 | 1.50E-07 | 1.30E-06 |
| | hippocampal fissure | 0.53 | 7.70E-09 | 1.20E-07 |
| | hippocampal tail | 0.6 | 4.50E-10 | 1.90E-08 |
| amygdala | accessory basal nucleus | 0.24 | 7.90E-03 | 0.023 |
| | paralaminar nucleus | 0.27 | 1.80E-03 | 0.0056 |
| | basal nucleus | 0.32 | 2.20E-04 | 0.0008 |

**Table S6. Gene dosage effects controlling for antipsychotic medication.** Repeat of gene dosage volume analysis with the addition of a covariate coding whether or not each participant was taking antipsychotic medication at the time of the scan. Regions with FDR $q < 0.05$ are listed in this table.
